# Pancreatic δ-cells are resistant to auto- and paracrine inhibition in human type-2 diabetes

**DOI:** 10.64898/2025.12.12.693197

**Authors:** Muhmmad Omar-Hmeadi, Lina Matuseviciene, Liangwen Liu, Per-Eric Lund, Sebastian Barg

## Abstract

Somatostatin secretion from pancreatic δ-cells inhibits nearby α-and β-cells, and tunes the body’s glycemic set-point. The role of δ-cells in diabetes remains unclear, in part due to the difficulty separating intrinsic regulation from intra-islet paracrine effects. Here we compared the function of isolated δ-cells of cadaveric non-diabetic and type-2 diabetic donors, by single cell TIRF-microscopy and electrophysiology. Elevated glucose stimulated exocytosis of somatostatin, which was further amplified by glucagon, exendin-4, or forskolin, independent of diabetic status. GABA enhanced exocytosis and electrical activity, while insulin had no effect. Adrenaline and somatostatin strongly inhibited δ-cell activity, leading to autocrine feedback inhibition of somatostatin exocytosis. In type-2 diabetes, δ-cell inhibition by somatostatin and adrenaline was lost, together with a marked reduction in somatostatin receptor (SSTR2) surface expression. We further show that resistance to somatostatin leads to hyperactive δ-cells in type-2 diabetes, and propose that this mechanism contributes to defective blood glucose control.

## Introduction

Blood glucose in vertebrates is primarily controlled by islets of Langerhans, specialized endocrine micro-organs in the pancreas that secrete the hormones insulin, glucagon, and somatostatin. Glucose-dependent insulin release from islet β-cells promotes peripheral glucose uptake and storage, while glucagon released from α-cells leads to glycogen breakdown and gluconeogenesis. Somatostatin release from δ-cells inhibits both α-and β-cells ^1,2^, and thereby tunes islet hormone output to control the glycemic set-point in mice ^3^. In rodents, δ-cells are concentrated at the islet periphery ^4–6^, while in humans δ-cells are scattered throughout the islet, with elongated filopodia-like structures that are in direct contact with α-and β-cells within the islet ^7^. The δ-cells secrete somatostatin-14 (SST-14), a 14-amino acid peptide that acts mostly paracrine and contributes minimally to circulating SST levels. SST activates Gαi-coupled somatostatin receptors (SSTRs) in α-and β-cells, which inhibits hormone release by a decrease in cytosolic cAMP and by activation of GIRK channels to dampen electrical activity ^1^. Secretion of somatostatin from isolated mouse islets is stimulated by glucose above 3 mM ^8^ and released in pulses that are synchronized with insulin secretion ^9^. As in β-cells, glucose controls δ-cells by closure of ATP-sensitive potassium (KATP) channels, action potential firing, and Ca^2+^-influx to trigger exocytosis of somatostatin-containing granules ^10,11^. Synchronization of δ-and β-cell activity is enhanced by electrical coupling via gap junctions ^12^. In addition, glucose stimulates somatostatin secretion by promoting cAMP-and Ca^2+^-dependent release of Ca^2+^ (CICR) from the endoplasmic reticulum ^13^. This pathway is particularly important during hyperglycemia, and is activated by a rise in intracellular Na^+^ ^14^ and limited by the ER resident two-pore K^+^ channel TALK-1 ^15^. Exocytosis of somatostatin granules is similar to that in other endocrine cells, where a series of preparatory steps results in release-ready (primed) granules that can be triggered to undergo exocytosis in response to influx of Ca^2+^ ^16^. In α-and β-cells, disturbances in the exocytosis machinery contribute to impaired hormone release in human type-2 diabetes ^2,17,18^.

Somatostatin secretion is modulated by hormones and neurotransmitters ^1^, consistent with the expression of the corresponding receptors in mouse δ-cells ^19,20^. Release of GABA from β-cells and of glutamate from α-cells both stimulate somatostatin secretion by activating ionotropic receptors and membrane depolarization ^21–23^, and it is plausible that this mechanism acts as a local negative feedback to limit release of insulin and glucagon. Somatostatin secretion is enhanced by the peptide hormones glucagon ^24^, GLP-1 ^25^, ghrelin ^19,20^, and urocortin-3 ^26^, likely via increased cytosolic cAMP. Adrenaline inhibits somatostatin secretion from intact islets, likely by activating the α2 adrenergic receptor ^20,27^. Insulin has been reported to inhibit, as well as stimulate somatostatin release ^19,27,28^. Interestingly, δ-cells also express receptors for somatostatin ^20^, suggesting negative autoinhibitory feedback. Modulation of δ-cell activity is expected to indirectly affect secretion of insulin and glucagon, and thus contribute to glucose homeostasis.

There is growing evidence for dysregulated somatostatin signaling in diabetes. Hypersecretion of somatostatin is seen in islets of human donors diagnosed with type-2 diabetes (T2D), and in high-fat diet fed (HFD) mice, a model of the disease ^7,29–31^. The δ-cells also undergo morphological changes during the pre-diabetic stage ^7^, and their number is reduced in T2D islets ^32^, suggesting early involvement of δ-cell dysfunction in the pathogenesis of T2D. In addition, α-cells of human T2D show a marked resistance to somatostatin that causes excessive glucagon release during hyperglycemia ^2^. In type-1 diabetes (T1D), the counter-regulatory glucagon response during hypoglycemia is impaired ^30,33^, which increases the risk for dangerous hypoglycemic events. At least in rat models of the disease, this can be rectified by treatment with somatostatin receptor antagonists, suggesting that hypersecretion of somatostatin plays a role in the disease. Although the hypersecretion of somatostatin in T1D likely results from reduced δ-β cell electrical coupling ^30^, it is not understood in T2D.

Understanding human δ-cell physiology is complicated by the scarcity of islet tissue, and difficulties interpreting somatostatin release data in experimental situations where paracrine interactions are left intact. Here, we used live cell imaging and patch clamp electrophysiology to study the regulation of SST exocytosis in individual dispersed δ-cells from non-diabetic (ND) and type-2 diabetic human donors. We report that δ-cell exocytosis is intrinsically glucose dependent in the physiological glycemic range. Importantly, δ-cell function in T2D is no longer inhibited by somatostatin or adrenalin, together with reduced membrane expression of the somatostatin receptor SSTR2. We show that this lack of auto-inhibition by somatostatin leads to increased somatostatin exocytosis, thereby providing a plausible mechanism for hypersecretion in T2D.

## Results

### Exocytosis of somatostatin granules in human δ-cells

Docking and exocytosis of somatostatin granules at the plasma membrane was studied in dispersed islet preparations from 45 non-diabetic donors (ND; see Suppl Table 1 and 3) that all had glycated hemoglobin HbA1c values ≤6% (mean ± SEM: 5.43 ± 0.04%, Fig. S1A). To identify δ-cells and at the same time fluorescently label somatostatin granules, we transduced the cells with an adenoviral vector (Psst-NPY-EGFP or Psst-NPY-Neon) to drive expression of a fluorescent protein labeled granule marker from the somatostatin promoter (see methods). After culture for 26–48 h, δ-cells were imaged by total internal reflection (TIRF) microscopy, which selectively images fluorescence near the plasma membrane (exponential decay constant *τ* ∼ 0.1 µm). Cells expressing the marker showed punctate staining that strongly co-localized with anti-somatostatin immunostaining (Fig. 1A). Local application of elevated K^+^ (75 mM, replacing Na^+^) to depolarize individual cells resulted in exocytosis, seen as rapid disappearance of individual fluorescently labeled granules (Fig. 1B, example in Fig 1C). We compared the rate of K^+^-stimulated exocytosis in cells bathed in either 10 or 1 mM glucose, in presence of the KATP channel opener diazoxide to prevent spontaneous depolarization. In 10 mM glucose, a 40-second long depolarization with elevated K^+^ (75 mM, replacing Na^+^) evoked exocytosis of 0.11 ± 0.01 granules μm^-^² (n = 84 cells from 11 donors; Fig. 1D). Exocytosis in 1 mM glucose was 0.052 ± 0.01 granules μm^-^²s^-1^, or about half of that in 10 mM glucose (n = 37 cells from 7 donors; p=0.003, Tukey post hoc test, Fig 1D). In both conditions, the time course of exocytosis suggested two kinetic components: a fast release phase lasting <10 s (7.2 ± 1× 10⁻³ granules μm^-^²s^-1^ during the first 10 s in 10 G), followed by a slower sustained phase (1.2± 0.2× 10⁻³ granules μm^-^²s^-1^). The rapid initial phase is consistent with depletion of a readily releasable pool (RRP) of granules, as previously described in β-and α-cells. Exocytosis occurred exclusively from granules that were immobile and present at the release site (docked) at the beginning of the recording. Docked granules, quantified as granule density visible by TIRF imaging prior to stimulation, was 0.72 ± 0.03 granules μm^-^² in 10 mM glucose, and 0.5± 0.03 granules μm^-^² in 1 mM glucose (n = 84 and 37 cells from 11 and 7 donors; Fig. 1E). In summary, somatostatin secretion in human δ-cells is mediated by exocytosis of docked somatostatin granules, elevated glucose primes these granules for exocytosis, and sustained stimulation partially depletes the docked granule pool.

**Fig 1.**
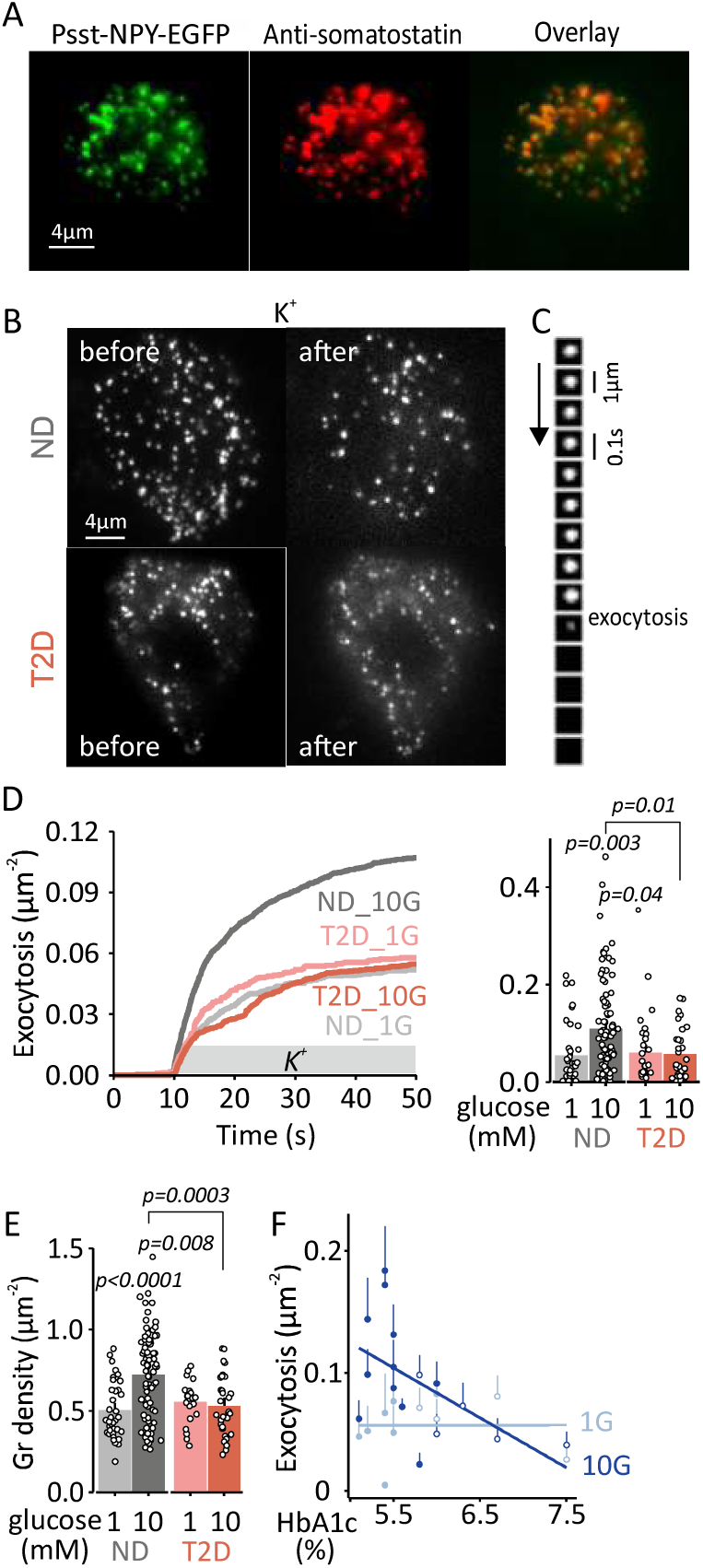
Loss of glucose dependent potentiation of δ-cell exocytosis in type-2 diabetes. A. Representative TIRF image of a human δ-cell transduced with *Psst*-NPY-EGFP (green, left) and immunostained for somatostatin (red, middle). Scale bar, 4 μm. B. Representative TIRF images of δ-cells from non-diabetic (ND) and type 2 diabetic (T2D) donors before and after stimulation with 75 mM K⁺, applied from 10 to 50 s. Diazoxide (100 µM) was present to prevent spontaneous glucose-dependent exocytosis. Scale bar, 4 μm. C. Image sequence showing exocytosis of a single granule in an experiment as in C. D. Time course of cumulative exocytotic events (left) and total exocytosis (right) during 75 mM K⁺ stimulation (10–50 s), in δ-cells from ND (gray) and T2D (red) donors, for glucose concentrations as indicated. Data are shown as mean ± SEM normalized to cell area; dots represent individual cells. ND 10G n = 84 cells, 11 donors; ND 1G n = 37 cells, 7 donors; T2D 10G n = 31 cells, 5 donors; T2D 1G n = 23 cells, 4 donors. Tukey post hoc: ND 10G > ND 1G (*p* = 0.003), ND 10G > T2D 10G (*p* = 0.011), ND 10G > T2D 1G (*p* = 0.044); all other comparisons not significant. E. Docked granule density (granules/μm²) for cells in D before stimulation (mean ± SEM). Tukey post hoc: ND 10G > ND 1G (*p* < 0.0001), ND 10G > T2D 10G (*p* = 0.0003), ND 10G > T2D 1G (*p* = 0.0076); all other comparisons not significant. F. Exocytosis (mean± SEM) as function of donor HbA1c at 1 mM (light blue) and 10 mM (dark blue) glucose. Lines are linear fits to the data, indicating difference between 1 and 10mM glucose (Pearsons correlation p=0.026, r=-0.55 for 10mMLinear regression model (R^2^=0.40),p=0,996 for HbA1c, p=0.036 for the glucose effect, and 0.062 for the HbA1c × glucose interaction.

### Reduced granule docking and exocytosis in δ-cells of T2D donors

We performed similar experiments with δ-cells from 18 donors that were diagnosed with type 2 diabetes (T2D; see Suppl Table 2 and 3) or had HbA1c values >6.0% (mean ± SEM: 6.9 ± 0.2%; Fig S1A). Stimulation with elevated K^+^ (in 10 mM glucose with diazoxide) led to exocytosis of 0.055 ± 0.010 granules μm^-^²s^-1^(n = 31 cells from 5 donors), corresponding to about half of that in ND cells (49% ± 13%, p = 0.01, Tukey post hoc). This reduction was primarily due to fewer primed granules compared with ND cells, as suggested by the fact that the initial fast phase was reduced to 2.8 × 10⁻³ granules μm^-^²s^-1^(p = 0.0003), while there was little change in the sustained phase (0.9 × 10⁻³ granules μm^-^²s^-^1; p = 0.3). Docked granule density in unstimulated T2D δ-cells was reduced by 27±6% compared with ND cells (0.53 ± 0.03 granules/μm² (n = 31 cells from 5 donors, p = 0.0003; Fig. 1E). We assessed the relationship between granule exocytosis and donor glycemic status (HbA1c). Pearson’s correlation analysis between HbA1c and K^+^ stimulated exocytosis revealed negative correlation (r =-0.55, p=0.026 for 10 G); linear regression analysis indicated that this was due to a loss of glucose dependence in T2D (Fig 1F). Exocytosis was also measured by patch clamp electrophysiology, as single cell capacitance increase in response to a train of 14 depolarizations (200 ms from –70 to 0 mV). The cumulative exocytotic response (Σ1–14 depolarizations) showed a trend towards reduced exocytosis in T2D δ-cells (24.6 ± 5 fF/pF, *n* = 33), compared with ND cells (40.3 ± 6 fF/pF, *n* = 45; *p* = 0.055, unpaired t-test; Fig. 2A-B). The initial whole-cell capacitance as measure of total cell surface area did not differ between ND and T2D δ-cells (9.1 ± 0.5 n = 33 vs 8.8 ± 0.6 pF n=45; Fig. 2C). The differences in exocytosis were not explained by altered activity of voltage-gated Na⁺ and Ca²⁺ channels, with similar peak inward currents and current–voltage relationships in ND and T2D δ-cells (Fig. 2D–F), indicating that the reduced exocytosis in T2D cells occurs downstream of depolarization and Ca²⁺ entry, likely at the level of granule docking, priming, or fusion.

**Fig 2.**
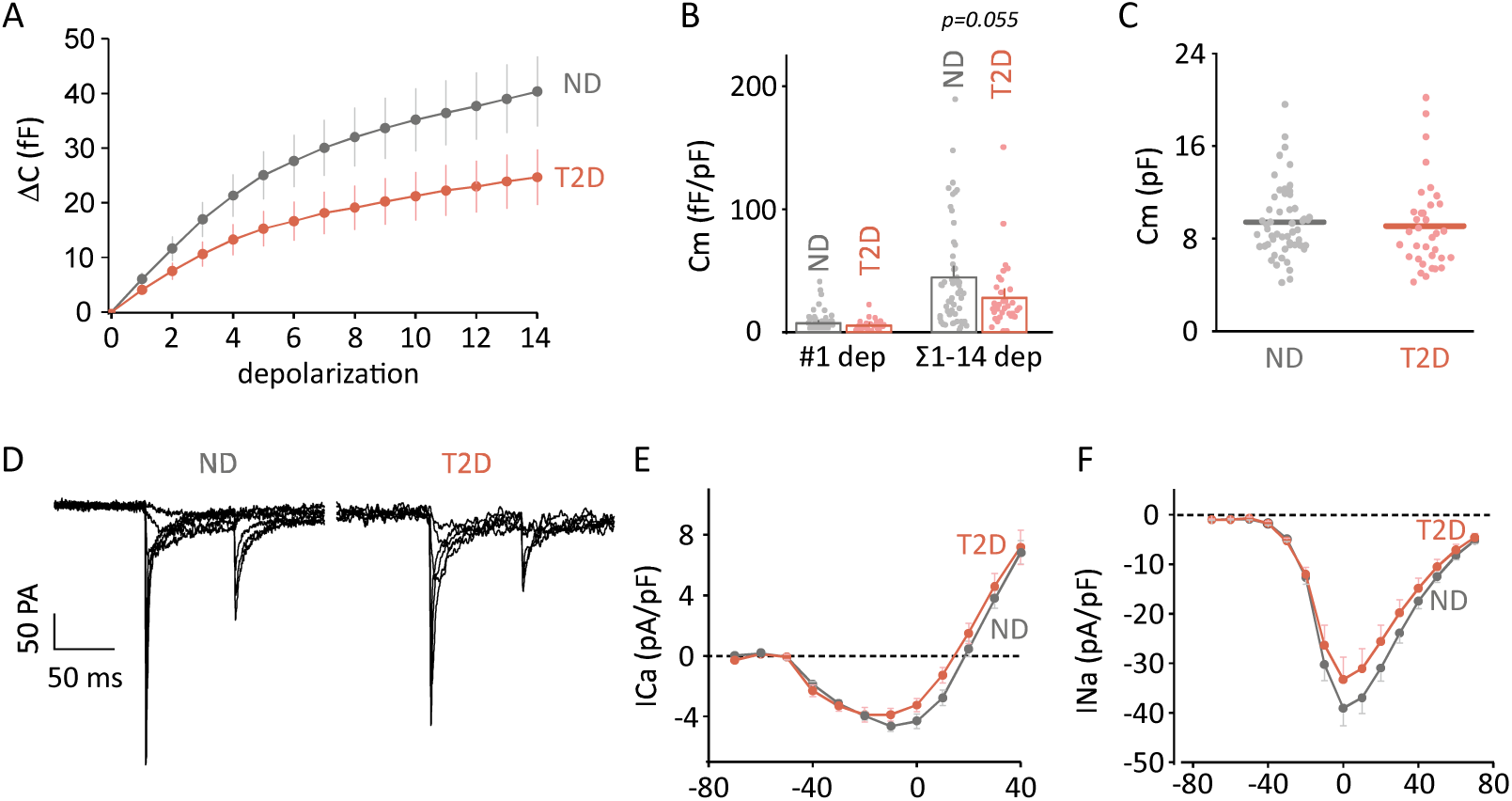
Voltage-dependent currents and exocytosis in human δ-cells Cumulative membrane capacitance increase (ΔCm) during a train of 14 depolarizing pulses (200 ms each, from-70 mV to 0 mV) in δ-cells from ND (grey) and T2D (red) donors; mean ± SEM. A. Membrane capacitance responses normalized to cell size (initial Cm), for the first depolarization (#1) and the total response (Σ1–14). Data represent mean ± SEM from ND (n = 45 cells, 11 donors) and T2D (n = 33 cells, 5 donors). B. Whole-cell membrane capacitance (Cm) in δ-cells from ND and T2D donors. Dots represent individual cells; lines indicate mean values. C. Representative families of voltage-dependent whole-cell currents recorded in δ-cells from ND and T2D donors. Currents were elicited by 50 ms depolarizations from –70 mV to +80 mV in 10 mV steps from a holding potential of –70 mV. D. Current (I)-voltage (V) relationship for Ca²⁺ currents, measured as average current during 5–45 ms of the depolarization shown in (D), normalized to cell capacitance (pA/pF). ND: n = 45 cells, 11 donors (gray); T2D: n = 31 cells, 5 donors (red). Mean ± SEM. E. I–V relationship for Na⁺ currents, measured as peak inward current during the first 5 ms of the depolarization shown in (D), normalized to cell capacitance (pA/pF). ND: n = 45 cells, 11 donors (grey); T2D: n = 31 cells, 5 donors (red). Mean ± SEM.

### Glucose regulation of somatostatin exocytosis

To investigate how physiological glucose concentrations regulate somatostatin exocytosis, we quantified spontaneous exocytosis (no diazoxide) in dispersed human δ-cells from ND and T2D donors at 1, 3, 7, and 10 mM glucose. Cells were pre-incubated in the respective glucose concentrations for at least 20 min, followed by a 3-minute recording period during which exocytosis was quantified (Fig. 3A). Spontaneous granule exocytosis was detected in all conditions, with a gradual increase in exocytosis frequency as glucose increased. Exocytosis rates (mean ± SEM, granules*μm^-^² per 3 min, Fig. 3B) increased linearly by more than three-fold within the tested glucose range (1 to 10 mM, *p*<0.0001 for glucose, n.s. for diabetes status, Two-way ANOVA). The density of docked granules tended to increase with glucose for both ND and T2D δ-cells (p=0.02 for ND and 0.07 for T2D, Fig. 3C). Values in ND cells were somewhat lower than in T2D and those in Fig1E, likely due to ongoing spontaneous exocytosis that depleted docked granules. To quantify the timecourse by which glucose promotes granule docking, we again prevented spontaneous exocytosis with diazoxide and recorded ND δ-cells during a shift from 1 to 10 mM glucose. Granule density increased from 0.55±0.02 to 0.75±0.02 granules*μm^-^², with a time constant of τ=11.5±4.7 minutes (p=7.4*10^-7^, Fig. 3D). We conclude that glucose augments somatostatin exocytosis by increasing the number of docked and primed granules, in addition to its effect on electrical activity. The potentiating effect of glucose is compromised in T2D.

**Fig 3.**
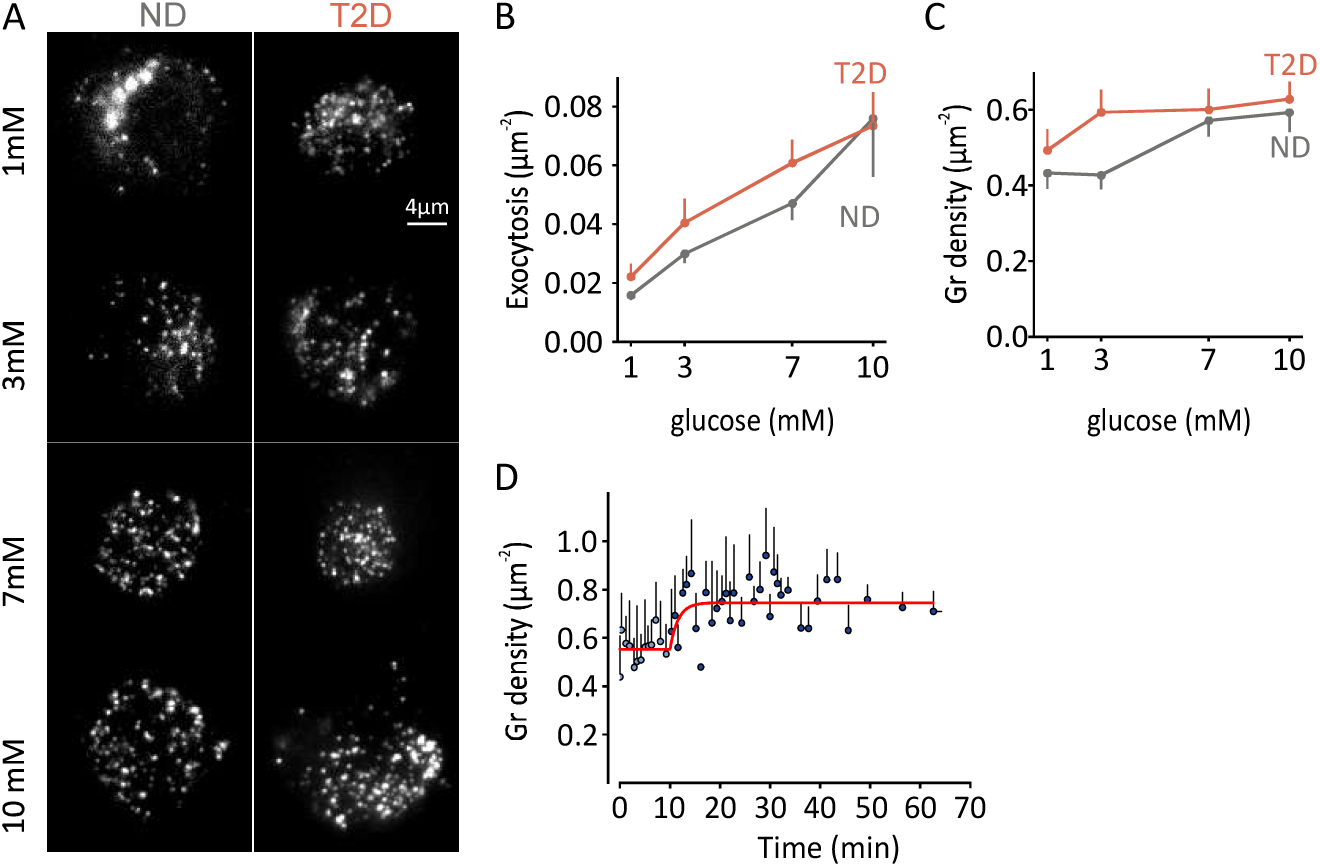
Glucose dependence of δ-cell exocytosis. A. Representative TIRF images of dispersed δ-cells from ND and T2D donors expressing Psst-NPY-Neon, exposed to the indicated glucose concentrations; no diazoxide was present. Scale bar, 4 µm. B. Quantification of spontaneous glucose-dependent exocytosis (mean ± SEM) during 3-minute recordings for cells as in A; dots represent individual cells. Effects of glucose are significant (*p* < 0.0001, two-way ANOVA and Tukey’s test). C. Granule density prior to stimulation at each glucose level for cells in B (mean ± SEM). D. Time course of granule density in ND δ-cells following glucose shift from 1 to 10 mM in presence of diazoxide. The line is a fitted a mono-exponential function (τ = 11.5± 4.7 min) to the data.

### Autocrine inhibition of exocytosis in dispersed δ-cells

Since human δ-cells express somatostatin receptor 2 (SSTR2), we tested for the effect of SST on exocytosis and membrane potential. In ND cells, somatostatin (SST, 400 nM) inhibited K⁺-stimulated exocytosis by 63% in 10mM glucose (p=0.0001 Fig 4A,C) and by 39% in 1mM glucose (p=0.0002; Fig 4B-C). In contrast, there was no effect of SST on exocytosis in T2D cells, regardless of glucose concentration. The loss of inhibition by SST was significant at high glucose concentrations (p= 0.0013 at 10mM and p= 0.76 at 1mM for ND vs T2D; Fig 4A-C). SST inhibition correlated with the donor’s glycemic status, HbA1c, at 1mM glucose (Pearsons p=0.083, r=0.69), but not at 10mM glucose (Pearsons p=0.58, r=0.18; Fig. 4D). SST had no effect on granule docking in any of the conditions (Fig. S2).

**Fig 4.**
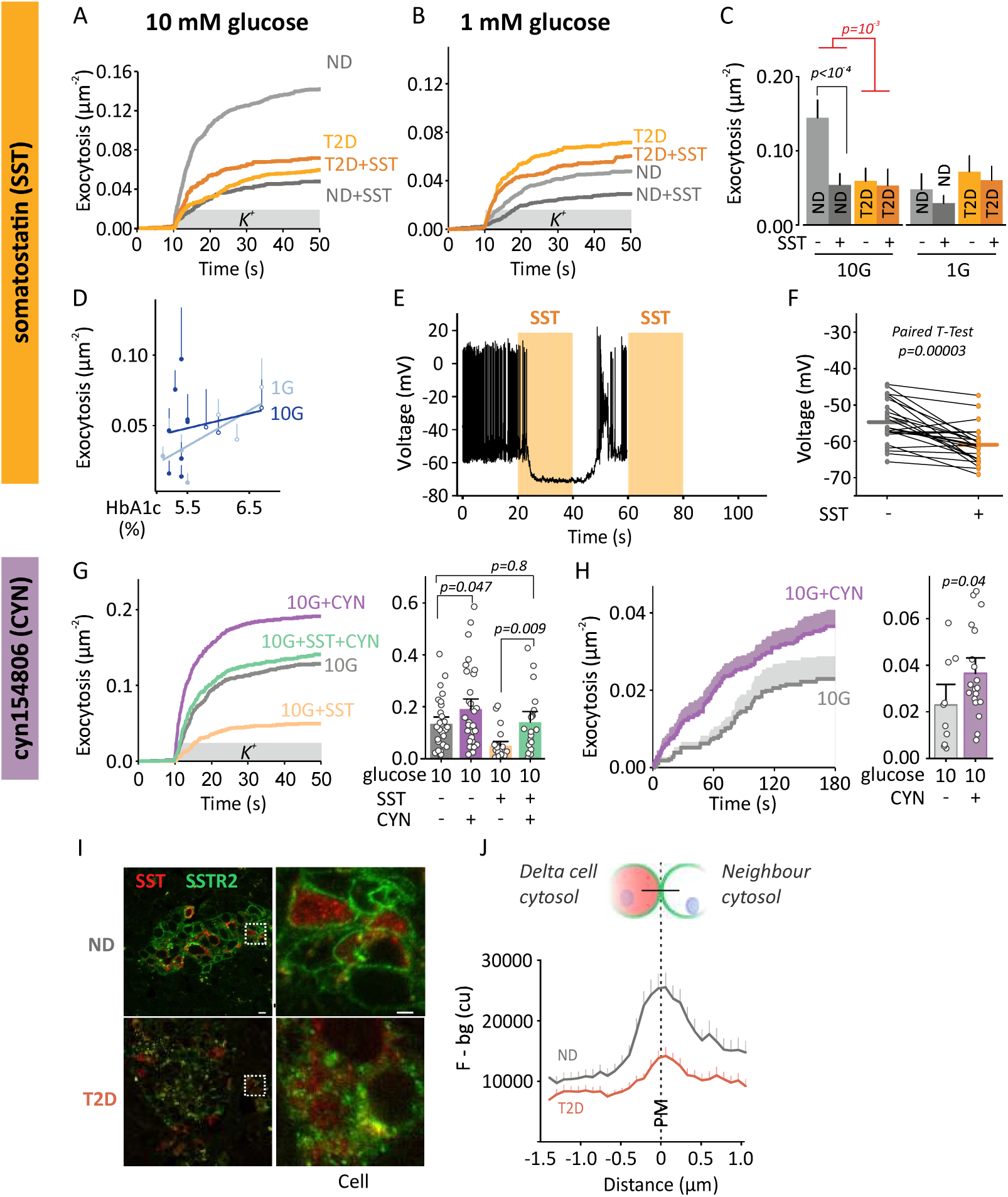
Reduced inhibition of δ-cell exocytosis by somatostatin in type-2 diabetes. A. Effect of somatostatin (SST, 400nM) on cumulative K⁺-stimulated exocytosis (mean±SEM) of ND (grey) and T2D (orange) δ-cells at 10 mM glucose (with diazoxide). 10G ND n= 45 cells, 6 donors; 10G ND +SST: n= 55 cells, 8 donors; 10G T2D no SST n= 24 cells, 4 donors; 10G T2D +SST n= 19 cells, 3 donors. B. As in (A), but at 1 mM glucose (1G). 1G ND no SST n= 26 cells, 5 donors; 1G ND +SST n= 21 cells, 4 donors; 1G T2D no SST n= 16 cells, 3 donors; 1G T2D +SST n= 21 cells, 3 donors. C. Average total exocytosis (granules/μm², mean ± SEM) in A and B. Significant effects of SST are indicated (three way ANOVA, post hoc Tukey test). D. Correlation between exocytosis and donor HbA1c at 1 mM (light blue) or 10 mM glucose (dark blue). Pearson’s p=0.08, r=0.69 for 1mM; p=0.2, r=-0.46 for 10mM glucose. E. Membrane potential recording from a representative ND δ-cell in 3 mM glucose. SST (400 nM) was applied during the orange-shaded intervals. F. Effect of SST on average membrane potential (mean±SEM) in ND δ-cells (25 cells/5 donors); 10 mM glucose. G. Cumulative (left) and total (right) K⁺ stimulated exocytosis (10 G with diazoxide) in presence of SST, SSTR2 antagonist (CYN154806, 200 nM), or both. 10 G: 36 cells/4 donors, 10 G + SST: 37 cells/4 donors, 10 G + CYN: 33 cells/3 donors, 10 G + SST + CYN: 33 cells/4 donors. Significant differences are indicated (two-tailed t-test). H. Cumulative (left) and total (right) spontaneous exocytosis (10 mM glucose, no diazoxide) during 3-minute recordings in presence (purple) or absence (grey) of CYN154806. 10G: 10 cells, 10G + CYN: 19 cells. I. Confocal images of human pancreatic tissue sections from ND (top) and T2D donors (bottom), stained for SSTR2 (green) and somatostatin (red). Scale bar: 10 µm. Right images magnify the boxed areas (SSTR2; scale bar: 2 µm). J. Fluorescence intensity profiles for SSTR2 staining in images as in I (mean±SEM), after subtraction of background fluorescence. Measurements were taken along lines placed perpendicular to the plasma membrane, as indicated schematically (ND: 289 profiles from 61 islet of 12 donors; T2D: 300 profiles from 41 islets of 7 donors). Average δ-cell fluorescence (-35%, p=0.0004, t-test) and the ratio of PM/δ-cell cytosol staining (-51%, p=0.0013, t-test) were different.

Next, we tested the effect of SST on membrane potential by perforated patch current-clamp recordings of dispersed δ-cells in 3 mM glucose without diazoxide (Fig. 4E, F). ND δ-cells displayed action potential firing, which was rapidly suppressed by acute application of SST (400 nM) from a pressurized capillary. Electrical activity typically resumed∼10 seconds after SST washout. On average, SST lowered the membrane potential by 6.2±1.6 mV (p = 0.00003), and inhibited action potentials in 80% of the cells (25 cells/5 donors; Fig. 4F). In some cases, activity re-emerged during SST application (Fig. S4), suggesting receptor desensitization or inactivation.

Inhibition of δ-cells by SST is expected to result in autocrine inhibition. We tested this notion by quantifying the effect of the SSTR2 antagonist CYN154806 (200 nM) on K^+^-stimulated exocytosis. As expected, the inhibition of exocytosis by SST was prevented in presence of CYN154806 (−55% vs −1.1%, p= 0.0005; Fig 4G). Importantly, CYN154806 increased K^+^-stimulated exocytosis also in absence of supplied SST (P=0.047, suggesting the presence of endogenous secreted SST. Likewise, glucose-induced exocytosis (10 mM, no diazoxide) was elevated by 60% (p=0.04) when CYN154806 was present (Fig 4H).

To understand the mechanism for reduced SST sensitivity in T2D δ-cells, we compared expression and the subcellular distribution of SSTR2 by immunostaining in pancreatic tissue sections of human ND and T2D donors (Fig. 4I). In ND islets, SSTR2 was detected in somatostatin-positive δ-cells and other islet cells, and was predominantly localized to the plasma membrane. In contrast, SSTR2 staining in T2D islets appeared markedly more diffuse, with reduced membrane staining. Quantitative analysis revealed a ∼50 % decrease in membrane-localized SSTR2 in T2D vs ND cell (Fig. 4J), despite similar somatostatin levels and distribution in both groups. Taken together, the data indicate that released SST mediates autocrine inhibition of δ-cell exocytosis and electrical activity. This auto-inhibition is lost in T2D, due to redistribution of SSTR2 away from the plasma membrane.

### β-cell derived signals: Insulin and GABA

Next, we examined the effect of the β-cell hormone insulin (100 nM). In ND δ-cells, addition of insulin had no significant effect on K⁺-evoked exocytosis or docked granule density (n.s, Tukey posthoc test), Fig. 5A, C, S2). In contrast, the β-cell transmitter molecule GABA increased exocytosis across all groups, with the greatest responses observed in ND cells. At 10 mM glucose, presence of GABA (400 nM) in the medium increased exocytosis in ND δ-cells 4.5-fold (p < 0.0001, Fig. 5D, F), and in T2D cells 3-fold (p = 0.009, Fig. 5D, F). In 1 mM glucose, GABA increased exocytosis of ND cells 3.3-fold (p=0.001), and exocytosis of T2D cells 3.8-fold (p = 0.0002, Fig. 5E, F). Granule docking was doubled by GABA in ND cells in 1 mM glucose (p = 0.001); this effect was reduced to 25% in T2D (p=0.006), but it was not seen at 10 mM glucose in either ND or T2D cells (Fig S2). GABA (1 µM, acutely applied) also caused near instantaneous membrane depolarization that ceased within 5–10 s after washout; GABA increased action potential frequency (p = 0.001, 40 cells from 5 ND donors, 3 mM glucose/no diazoxide, Fig. 5G-H). Given the stronger response to GABA compared with insulin, it is likely that the effect of GABA release from β-cells dominates and will promote δ-cell exocytosis and somatostatin release in intact islets.

**Fig 5.**
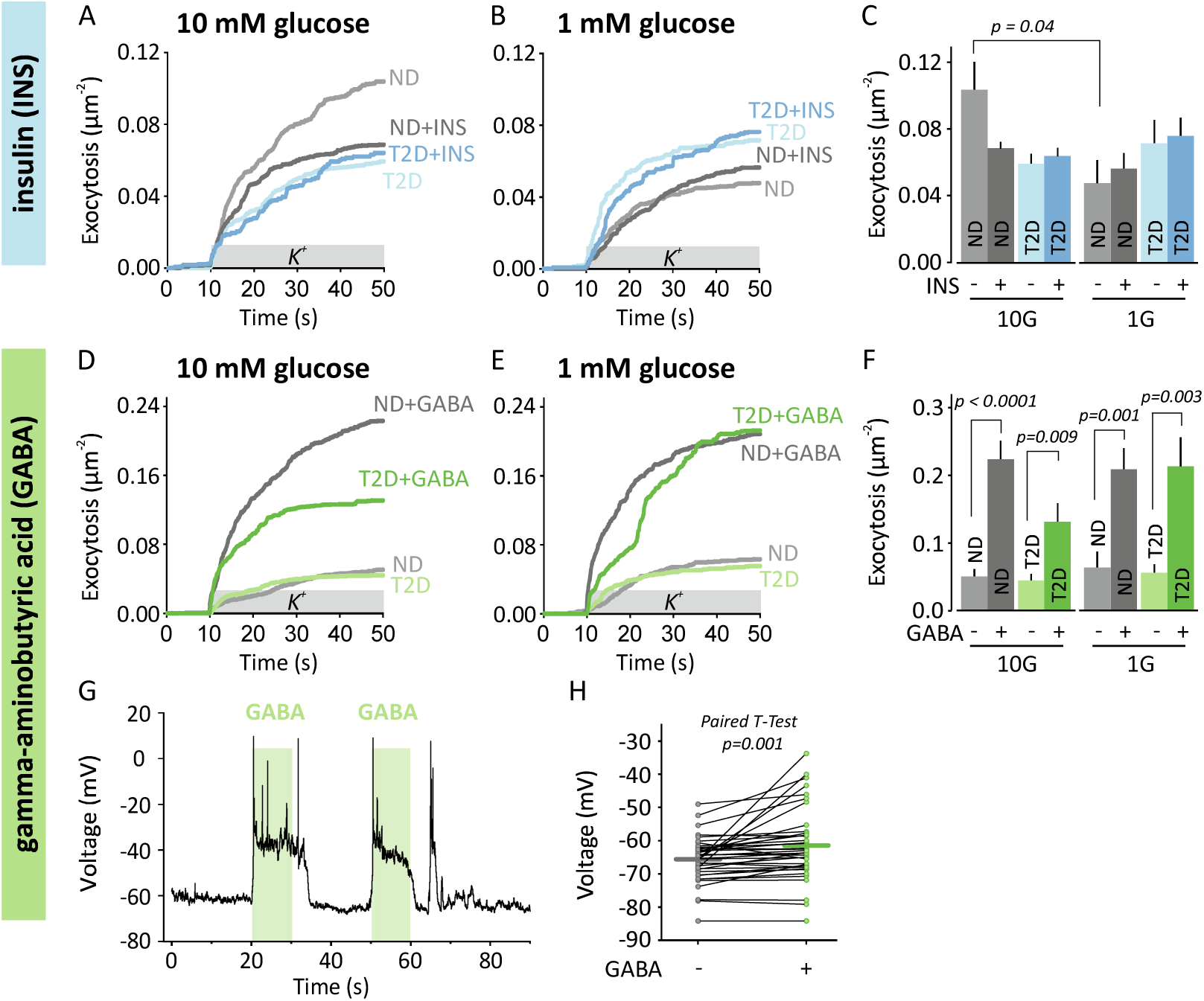
Effects of GABA and insulin on δ-cell exocytosis. A. Effect of insulin (100nM) on cumulative K⁺-stimulated exocytosis (mean±SEM) of ND (grey) and T2D (blue) δ-cells at 10 mM glucose (with diazoxide). ND control *n* = 18 cells, 3 donors; ND + insulin *n* = 24 cells, 4 donors; T2D control *n* = 14 cells, 3 donors; T2D + insulin *n* = 13 cells, 3 donors. B. As in A, but at 1 mM glucose. ND control *n* = 18 cells, 3 donors; ND + insulin *n* = 23 cells, 4 donors; T2D control *n* = 14 cells, 3 donors; T2D + insulin *n* = 13 cells, 3 donors. C. Average total exocytosis in A-B. Indicated comparison is significantly different (ANOVA, Tukey’s test). D. Effect of GABA (400nM) on cumulative K⁺-stimulated exocytosis (mean±SEM) of ND (grey) or T2D (green) δ-cells at 10 mM glucose. ND control n = 18 cells, 3 donors; ND + GABA n = 21 cells, 4 donors; T2D control n = 14 cells, 3 donors; T2D + GABA n = 17 cells, 3 donors. E. As in D, but at 1 mM glucose. ND control *n* = 11 cells, 2 donors; ND + GABA = 14 cells, 3 donors; T2D control *n* = 14 cells, 3 donors; T2D + GABA *n* = 13 cells, 3 donors. F. Average total exocytosis in D-E. Indicated comparisons are significantly different (ANOVA, Tukey’s test). G. Representative membrane potential recording from a dispersed ND δ-cell in 3 mM glucose. GABA (1 µM) was applied during the green-shaded interval. H. Average membrane potential before and after GABA application (1 µM) in ND δ-cells (40 cells/5 donors, p=0.001 paired t-test).

### Glucagon and cAMP signaling

We tested the effects of activating cAMP signaling on exocytosis in δ-cells of ND or T2D donors. K^+^-stimulated exocytosis was potentiated by glucagon (10 nM in the bath), with an about 2-fold increase in 10 mM glucose (p=0.02 and 0.06) and 3-fold in 1 mM glucose (p=0.0001 for both). There were no differences between ND and T2D groups (Fig 6A-C), and no correlation between glucagon’s stimulatory effect and donor HbA1c (Fig 6D). Linear regression model analysis confirmed that glucose affected exocytosis (p=0.014), independent of HbA1c p=0.6. Docked granule density nearly doubled in presence of glucagon in ND cells at 1 mM glucose (p = 0.002), and was slightly reduced (p = 0.0035) at 1mM glucose or without any effect (T2D at 1 or 10mM glucose). Similarly, the GLP-1 receptor agonist exendin-4 (EXN; 10 nM), and the adenylate cyclase activator forskolin (FSK; 2 µM) increased exocytosis 2 to 3-fold, with no effect of diabetes diagnosis or HbA1c (Fig 6E-L). These functional changes were accompanied by increased granule docking (Fig S2). In both ND and T2D δ-cells, docked granule density rose by about 50% in response to either of the three cAMP elevating agents, regardless of the glucose concentration (1 or 10 mM). We conclude that cAMP signaling in the δ-cell robustly potentiates somatostatin exocytosis.

**Fig 6.**
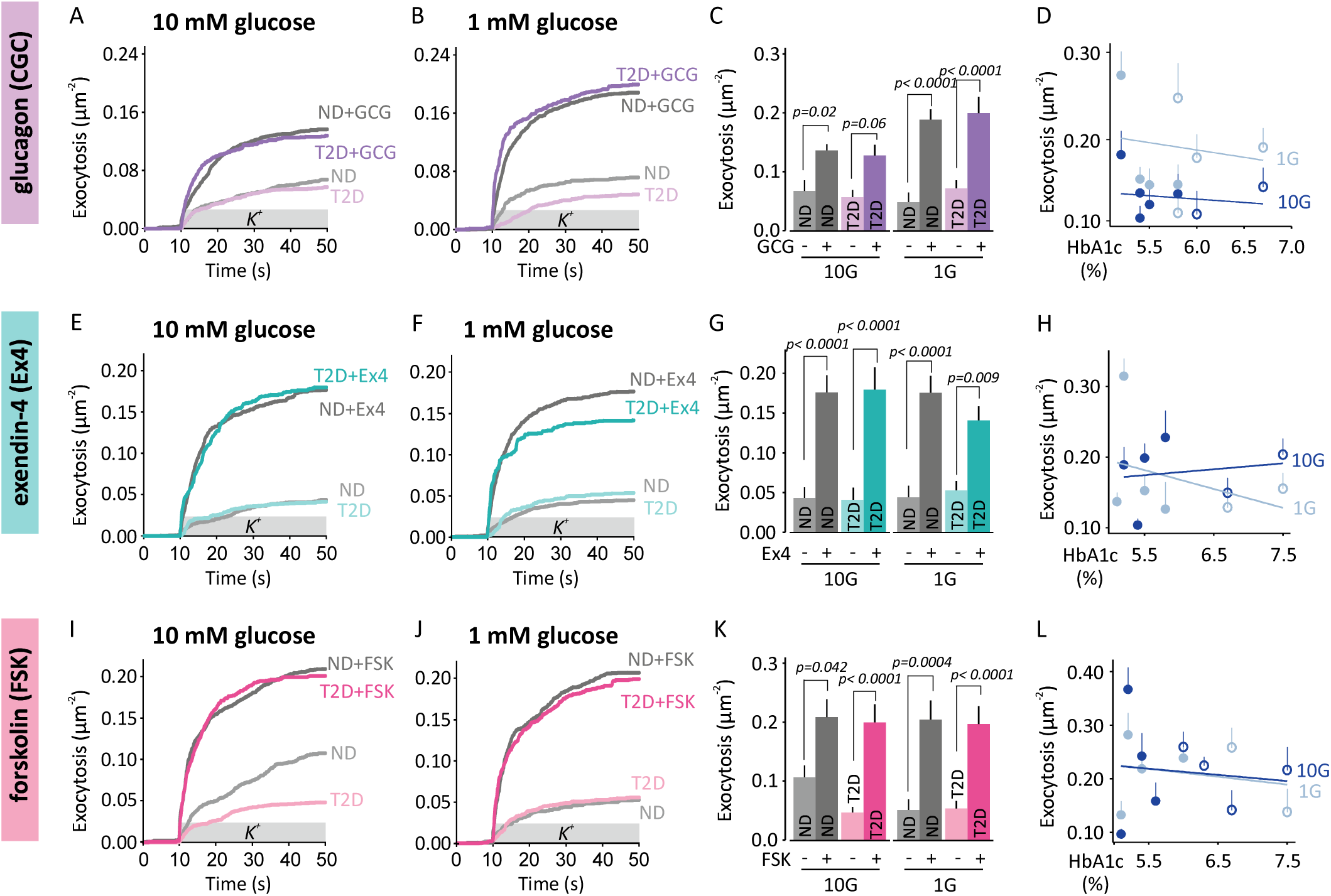
Glucagon, exendin-4, and forskolin amplify δ-cells exocytosis. A. Effect of glucagon (10nM) on cumulative K⁺-stimulated exocytosis (mean±SEM) of ND (grey) or T2D (purple) δ-cells at 10 mM glucose (with diazoxide). ND control *n* = 22 cells, 4 donors; ND + glucagon *n* = 28 cells, 4 donors; T2D control *n* = 20 cells, 3 donors; T2D + glucagon *n* = 17 cells, 3 donors. B. As in A, but at 1 mM glucose. ND control *n* = 19 cells, 3 donors; ND + glucagon *n* = 26 cells, 3 donors; T2D control *n* = 16 cells, 3 donors; T2D + glucagon *n* = 13 cells, 3 donors. C. Average total exocytosis in A-B. Indicated comparisons are significantly different (ANOVA, Tukey’s test). D. Correlation between granule exocytosis and donor HbA1c under 1 mM (light blue) and 10 mM (dark blue) glucose. Linear regression model yielding p = 0.603 for HbA1c and p = 0.014 for glucose with overall model significance of p = 0.042. E. Effect of exendin-4 (10 nM) on cumulative K⁺-stimulated exocytosis (mean±SEM) of ND (grey) and T2D (turquoise) δ-cells at 10 mM glucose. ND control *n* = 18 cells, 3 donors; ND + EXN *n* = 18 cells, 3 donors; T2D control *n* = 13 cells, 2 donors; T2D + EXN *n* = 9 cells, 2 donors. F. As in E, but at 1 mM glucose. ND control *n* = 22 cells, 3 donors; ND + EXN *n* = 19 cells, 3 donors; T2D control *n* = 14 cells, 2 donors; T2D + EXN *n* = 11 cells, 2 donors. G. Average total exocytosis in E-F. Indicated comparisons are significantly different (ANOVA, Tukey’s test). H. Correlation of exocytosis and donor HbA1c, as in D, for exendin-4 data in E-F. I. Effect of forskolin (2µM) on cumulative K⁺-stimulated exocytosis (mean±SEM) of dispersed ND (grey) and T2D (pink) δ-cells at 10 mM glucose. ND control n = 15 cells, 3 donors; ND + FSK n = 22 cells, 3 donors; T2D control n = 27 cells, 3 donors; T2D + FSK n = 17 cells, 3 donors. J. As in I, but at 1 mM glucose. ND control n = 16 cells, 3 donors; ND + FSK n = 16 cells, 3 donors; T2D control n = 19 cells, 3 donors; T2D + FSK n = 13 cells, 3 donors. K. Average total exocytosis in i-J. Indicated comparisons are significantly different (ANOVA, Tukey’s test). L. Correlation between exocytosis and donor HbA1c, as in D, for forskolin data in I-J.

### Adrenaline mediated inhibition is altered in T2D

To explore sympathetic regulation of δ-cell secretion, we tested adrenaline (5 µM, in the bath). In ND cells, adrenaline inhibited exocytosis, reducing K⁺-stimulated exocytosis by about 70% regardless of the glucose concentration (p=0.015 at 10mM glucose, p=0.003 at 1mM). Surprisingly, this effect was reversed in T2D δ-cells and adrenaline now doubled exocytosis at 1 mM glucose (p=0.01; Fig. 7B-C), but had no significant effect at 10 mM glucose (p=0.34, Fig 7A-C). This striking switch was apparent also when the fold-change in exocytosis (adrenaline/control) was plotted against donor HbA1c ( p = 0.002, r=0.9 at 1G; p = 0.008, r=0.77 at 10G; Fig. 7D). Granule density was not affected by adrenaline in any of the groups. Finally, membrane potential recordings showed that adrenaline rapidly suppressed δ-cell electrical activity in ND cells (3 mM glucose; Fig. 7E). This effect was consistent across donors (*n* = 13 cells, *p* = 0.04; Fig. 7F), and is expected to act in addition to the direct effect of adrenaline on exocytosis. We conclude that adrenaline robustly inhibits somatostatin exocytosis in ND δ-cells, and that it may lose effect or even become stimulatory in T2D.

**Fig 7.**
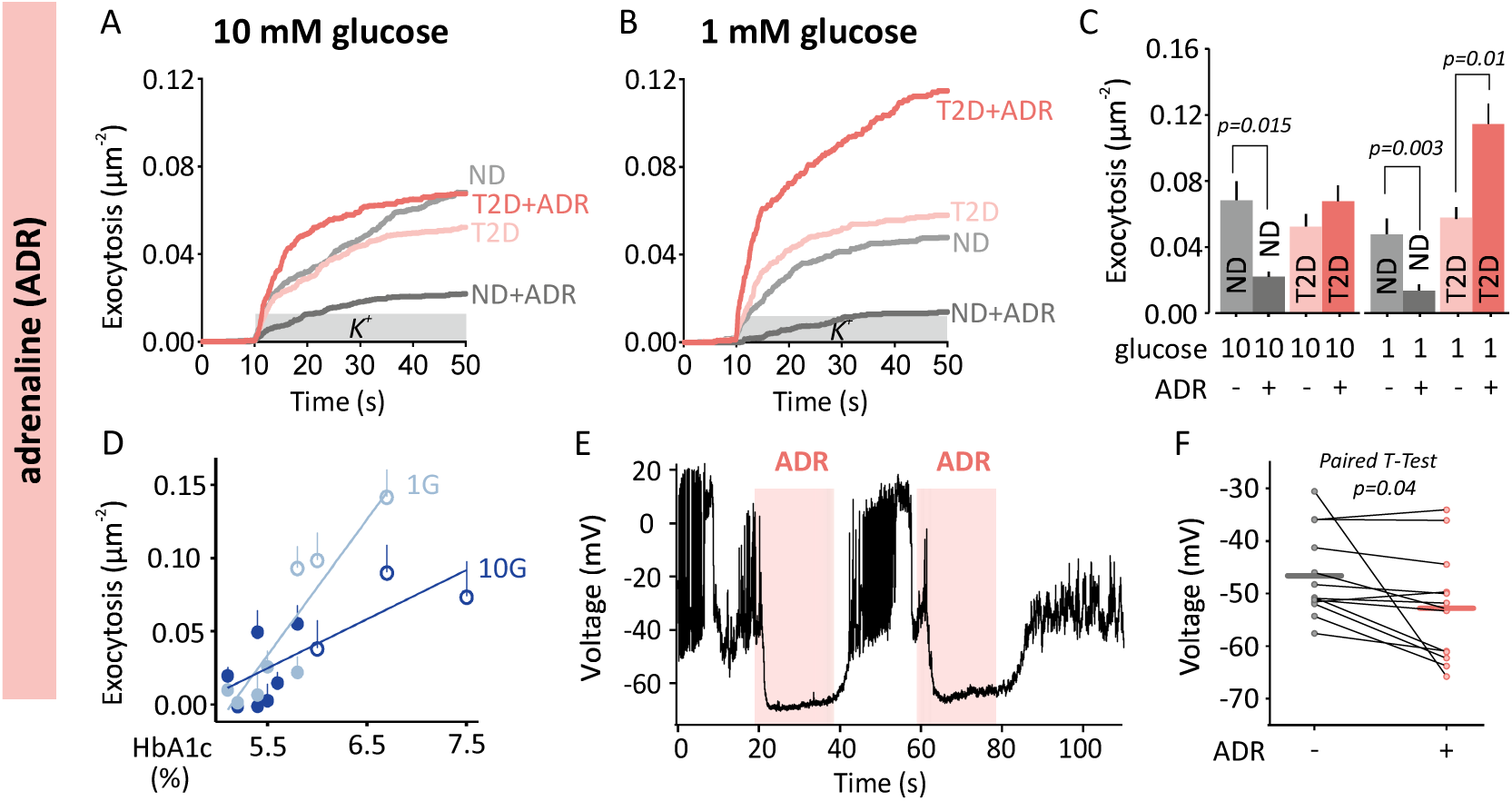
Adrenaline stimulates δ-cell exocytosis in type-2 diabetes. A. Effect of adrenaline (ADR, 5 µM) on cumulative K⁺-stimulated exocytosis (mean±SEM) of dispersed ND (grey) and T2D (red) δ-cells at 10 mM glucose (with diazoxide). ND control n = 28 cells, 4 donors; ND + ADR n = 36 cells, 4 donors; T2D control n = 27 cells, 3 donors; T2D + ADR n = 22 cells, 3 donors. B. As in A, but at 1 mM glucose. ND control n = 26 cells, 3 donors; ND + ADR n = 31 cells, 3 donors; T2D control n = 23 cells, 3 donors; T2D + ADR n = 15 cells, 3 donors. C. Average total exocytosis in A-B. Indicated comparisons are significantly different (ANOVA, Tukey’s test). D. Correlation of granule exocytosis and donor HbA1c at 1 mM (light blue) and 10 mM glucose (dark blue). Pearsons p=0.002, r=0.9 for 1mM; p=0.008, r=0.77 for 10mM glucose. Linear regression tested effects of HbA1c, glucose and their interaction on exocytosis: HbA1c p < 0.001; glucose p = 0.013; interaction p = 0.008; overall model p = 0.00012. E. Representative membrane potential recording from an ND δ-cell in 3mM glucose. Adrenaline (5 µM) was applied during the red-shaded interval. F. Average membrane potential in ND δ-cells following before and during adrenaline application (5 µM; n= 13 cells, *p* = 0.04 paired t-test).

## Discussion

Here we examined how paracrine and metabolic cues affect the function of single dispersed human δ-cells from non-diabetic (ND) and type 2 diabetic (T2D) islet donors. Our data suggest that key stimulatory pathways remain largely intact in T2D δ-cells, while important (auto)-inhibitory pathways are compromised. This selective loss of negative feedback may contribute to dysregulated somatostatin secretion and disturbed islet hormone balance and glycemic control in diabetes.

We show here that single human δ-cell function is intrinsically glucose sensitive, and exocytosis rates increased about fourfold as extracellular glucose rose from 1 to 10 mM. δ-cells are depolarized by glucose via the classical KATP-channel dependent triggering pathway, leading to action potential firing and Ca^2+^-influx ^10^. In addition, δ-cells are electrically coupled to β-cells through gap junctions ^12,34^, but our single cells data argue that this effect is not required for the glucose sensitivity of δ-cell exocytosis. There is also evidence that glucose can stimulate somatostatin secretion through a KATP channel independent mechanism, possibly through the operation of the glucose transporter SGLT1 ^19,20^. In addition, glucose potentiates exocytosis and granule docking independent of membrane potential depolarization (e.g. in presence of diazoxide), which indicates direct effects at the exocytosis machinery that lead to a rapid increase in the number of primed granules. Differences between ND and T2D δ-cells in glucose sensitivity were only minor compared with those we previously reported in β-cells ^2,17^.

Stimulation of cAMP signaling by glucagon, exendin-4, or forskolin accelerated somatostatin granule exocytosis several-fold, largely through an expanded readily releasable pool of granules, and with similar effect in in both ND and T2D δ-cells. Since glucagon receptor (GCGR) and glucagon-like peptide 1 receptor (GLP-1R) are expressed in δ-cells ^20^, the likely mechanism is through Gs-coupled elevation of cAMP and PKA activation as in β-cells. Notably, δ-cells of T2D donors remain responsive to these agents, which suggests that cAMP-dependent signaling to the exocytotic machinery is unaffected in the disease. Exocytosis of δ-cells was also stimulated by GABA, which is co-released with insulin from β-cells under hyperglycemic conditions. It is possible that GABA in this way acts as part of a β-δ-β feedback loop that limits excessive release of insulin (and co-released GABA) from β-cells. Since inhibitory SST signaling is lost in T2D, GABA may instead contribute to chronic hypersecretion of somatostatin. GABA application rapidly depolarized δ-cells, suggesting a mechanism where activation of ionotropic GABA-A receptors leads to Cl^-^ efflux. This is similar to β-cells, where high cytosolic [Cl^-^] results in a reversal potential of around-10 mV. In contrast, α-cells are hyperpolarized by GABA, reflecting lower cytosolic [Cl^−^] ^22^. Indeed, gap-junction coupling between β-and δ-cells, but not α-cells, is expected to equilibrate Cl^-^ of the connected cells.

In contrast, inhibitory pathways mediated by somatostatin and adrenaline were lost or even reversed in T2D δ-cells. We show that somatostatin exerts autocrine inhibition in ND cells, and both exocytosis and electrical activity were reduced substantially. This is consistent with SSTR2-mediated hyperpolarization (Fig 4E) and depriming of somatostatin granules (Fig 4C), which is similar to β-cells ^35^. In T2D δ-cells, somatostatin failed to inhibit exocytosis and action potential firing, in line with the reduced surface expression of SSTR2 in pancreas sections of T2D donors. It is likely that insensitivity to autocrine inhibition contributes to the increased exocytosis seen in T2D δ-cells. Adrenaline suppressed depolarization-evoked granule fusion, which is consistent with the expression of α2-adrenergic receptors in human δ-cells ^36^. Notably, in T2D δ-cells adrenaline reverted to being stimulatory at low-glucose conditions. If this finding reflects the in vivo situation, this mechanism would impair the counter-regulatory glucagon response during severe hypoglycemia. This is because the increase in SST response to adrenaline would in turn block α-cell activity and inhibit glucagon release. Indeed, there is evidence for such a similar mechanism in T1D^30^. Although Insulin has been proposed to contribute to β to δ-cell signaling, we failed to detect direct effects of insulin on exocytosis of isolated δ-cells. However, there is good evidence for stimulation of δ-cells by molecules that are co-released with insulin during β-cell activity, including urocortin-3 ^26^ and GABA (Fig 5).

Our findings suggest that selective loss of inhibitory control in δ-cells in T2D lead to elevated somatostatin levels within the islet. In vivo, this is increase in SST release will reduce glucose-dependent secretion of insulin and glucagon, and thereby impede adequate responses to both hyper-and hypoglycemia. Elevated somatostatin may further lead to somatostatin resistance and SSTR2 internalization in α-cells of T2D donors, which in turn causes improper glucagon release during hyperglycemia ^2^. Conversely, elevated somatostatin at low glucose may reduce glucagon secretion and thereby impair the counter-regulatory response to hypoglycemia, a common problem in both type-1 and type-2 diabetes that affects quality of life. Adrenaline, the body’s other acute response to hypoglycemia, normally prevents somatostatin release in this situation, which we confirm here. In T2D δ-cells, adrenalin instead stimulated somatostatin release, which suggests that adrenaline may indirectly impair the glucagon response during hypoglycemia. Restoring somatostatin receptor signaling or α2-adrenergic responsiveness in δ-cells may therefore represent a strategy to rebalance islet hormone secretion and improve glycemic control. Conversely, activation of cAMP-dependent pathways - through GLP-1 receptor agonists or GABA modulators - may offer a means to modulate somatostatin output without provoking dysregulated secretion.

## Methods

### Human δ-cells

Human pancreatic islets were obtained from the Nordic Network for Clinical Islet Transplantation Uppsala ^37^ and the ADI Isletcore at the University of Alberta ^38^ with approval by the Uppsala Regional Ethics Board (2006/348) and the Alberta Human Research Ethics Board (Pro00001754), respectively. Experiments with isolated islets are approved by Etikprövningsmyndigheten (Dnr 2024-07948-02). Written informed consent was obtained from all donor families (Suppl Tables 1-3). Islets were cultured in CMRL-1066 medium containing 5.5 mM glucose, 10% fetal calf serum, 2 mM L-glutamine, 100 U/mL penicillin, and 100 U/mL streptomycin at 37°C in a humidified atmosphere with 5% CO₂ for up to two weeks. For single-cell experiments, islets were dispersed using cell dissociation buffer (Thermo Fisher Scientific) supplemented with 0.005% trypsin (Life Technologies). Cells were then washed and plated onto 22-mm poly-L-lysine-coated coverslips in serum-containing medium and allowed to settle overnight before transduction. To simultaneously identify human δ-cells and label somatostatin granules, we transduced dispersed islet cells with adenoviral constructs expressing neuropeptide Y (NPY) fused to EGFP or Neon under control of the somatostatin promoter (Psst-NPY-EGFP or Psst-NPY-Neon). We validated the specificity of δ-cell labelling by immunostaining; 91% of Psst-NPY-EGFP-positive cells were also somatostatin-positive, and approximately 50% of all somatostatin-positive cells were labelled with Psst-NPY-EGFP (Fig 1A). Colocalization analysis revealed that 89 ± 2% of somatostatin-positive granules also contained NPY-EGFP (14 cells from 3 donors; Fig 1A).

### Immunostaining of dispersed islet cells and pancreatic tissue sections

Dispersed cells transduced with Psst-NPY-EGFP were fixed in 4% paraformaldehyde and permeabilized with Triton X-100. After blocking, cells were incubated overnight at 4°C with primary antibodies against somatostatin, insulin, or glucagon. Alexa Fluor® 488-conjugated secondary antibodies were applied for 2 hours in the dark. Stained cells were mounted in DAPI-containing medium and imaged by confocal microscopy. Human pancreatic tissue sections were obtained from the Uppsala biobank, and staining experiments are approved by Etikprövningsmyndigheten (Dnr 2024-07948-02). Sections were deparaffinized and rehydrated using xylene and graded ethanol. Antigen retrieval was performed by heating in 10 mM sodium citrate buffer (pH 6.0), followed by 30 min cooling and PBS washes. Sections were blocked in 5% BSA (in Dako 1× buffer) for 10 minutes and incubated overnight at 4°C with primary antibodies: anti-somatostatin (ThermoFisher, MA5-17182; 1:800) and anti-SSTR2 (Abcam, ab134152; 1:400). After washing, sections were incubated with Alexa Fluor® 488 and 568 secondary antibodies (1:500 in wash buffer) for 30 minutes. Nuclei were counterstained using DAPI-mounting medium.

### Microscopy

All live cell imaging was performed using a custom-built lens-type Total internal reflection fluorescence (TIRF) microscope based on a Zeiss AxioObserver Z1 equipped with a 100×/1.45 NA oil-immersion objective (Carl Zeiss). Excitation was provided by 491 nm and 561 nm DPSS lasers (Cobolt), controlled via an acousto-optical tunable filter (AA-Opto). Excitation and emission paths were separated using a multiband dichroic beamsplitter (ZT405/488/561/640rpc, Chroma). Emitted light was further split into two channels with an image splitter (Optical Insights) using a 565 nm dichroic (565dcxr, Chroma) and filtered through bandpass filters (ET525/50m and 600/50m, Chroma) before detection with an EMCCD camera (QuantEM 512SC, Roper Scientific). Image scaling was 160 nm/pixel. For all experiments, cells were bathed in a standard extracellular solution containing (in mM): 138 NaCl, 5.6 KCl, 1.2 MgCl₂, 2.6 CaCl₂, 10 D-glucose, and 5 HEPES (pH 7.4, adjusted with NaOH). Glucose concentrations were varied as indicated (Figs. 3–6). Cells were equilibrated for at least 20 minutes before imaging. For K⁺-induced depolarization, Na⁺ in the solution was replaced equimolarly with 75 mM KCl. High K⁺ was applied from 10–50 seconds using a computer-controlled pressure ejection pipette positioned near the cell. Spontaneous exocytosis under glucose stimulation (Fig. 3A,B) was recorded for 3 minutes following equilibration in the stated glucose concentration. Confocal microscopy was performed using a Zeiss LSM780 with a 63/1.40 objective (Zeiss). The red channel (excitation 561 nm, emission 578–696 nm) and green channel (excitation 488 nm, emission 493–574 nm) were sequentially scanned. The pinhole size was 0.61 mm, which corresponds to 1 Airy unit.

### Image analysis

Exocytosis events were identified as sudden losses of granule fluorescence within 1–2 frames, and docked granules were quantified using the’Find Maxima’ function in ImageJ (NIH). Both values were normalized to the cell footprint area. SSTR2 distribution was analyzed using ImageJ. Square ROIs (32 × 32 pixels) were placed on SST-positive, DAPI-labeled cells with the cell membrane centered in the square, avoiding the nucleus. ROIs were background-subtracted and aligned to produce a composite image. Fluorescence intensity along a 3-pixel-thick line across the membrane was measured as shown in Fig. 4J.

### Electrophysiology

Whole-cell voltage clamp and capacitance measurements were performed using an EPC-9 amplifier (HEKA Electronics) with PatchMaster software. Patch pipettes (2–4 MΩ) were pulled from borosilicate glass, coated with Sylgard near the tip, and fire-polished. Recordings were performed at 32°C. Intracellular solution contained (in mM): 125 Cs-glutamate, 10 CsCl, 10 NaCl, 1 MgCl₂, 0.05 EGTA, 3 Mg-ATP, 0.1 cAMP, and 5 HEPES (pH 7.2, adjusted with CsOH). The extracellular solution contained (in mM): 138 NaCl, 5.6 KCl, 1.2 MgCl₂, 2.6 CaCl₂, 3 D-glucose, and 5 HEPES (pH 7.4, adjusted with NaOH), superfused at 0.4 mL/min. Voltage-gated Na⁺ and Ca²⁺ currents were evoked by 50 ms depolarizing steps from −70 mV to +80 mV in 10 mV increments. Capacitive and leak currents were subtracted using a P/4 protocol. Na⁺ current was defined as the peak within 0–3 ms; sustained current (5–45 ms) was attributed to Ca²⁺ influx. Exocytosis was measured as changes in cell capacitance using the lock-in module (1 kHz sine wave, 30 mV peak-to-peak). For current-clamp recordings in Fig. 4, the internal solution contained (in mM): 76 K₂SO₄, 10 KCl, 1 MgCl₂, and 5 HEPES (pH 7.4, adjusted with KOH).

### Statistics

All quantitative data are presented as mean ± SEM unless otherwise indicated. Data were analyzed using either two way or three way ANOVA, depending on the experimental design. Two-way ANOVA was applied when two independent factors (e.g., donor diabetes status and glucose concentration) were considered. Three-way ANOVA was used when analysis included glucose concentration (1 mM or 10 mM), donor diabetes status (ND or T2D), and treatment (e.g., somatostatin, insulin, GABA, glucagon, exendin-4, forskolin, adrenaline) as fixed factors. Where significant main effects or interactions were detected, Tukey’s Honest Significant Difference (HSD) test was used for post hoc pairwise comparisons. Levene’s test was used to assess homogeneity of variances. For correlation analyses (e.g., between exocytosis and donor HbA1c), Pearson’s correlation coefficients (r) and linear regression were computed. Paired or unpaired two-tailed Student’s t-tests were used for membrane potential or immunofluorescence comparisons, as appropriate.

## Acknowledgements

We thank Jan Saras for expert technical assistance and help with adenovirus design and preparation. Human islets were provided by the Nordic Network for Islet Transplantation (JDRF award 31-2008-416, ECIT Islet for Basic Research Program, and EXODIAB), the Alberta Diabetes Institute Islet-Core with assistance of the Human Organ Procurement and Exchange [HOPE] program, Trillium Gift of Life Network [TGLN], and other Canadian organ procurement organizations. The work was supported by the Swedish Science Council, Diabetes Wellness Network Sweden, the Swedish Diabetes Society, Barndiabetesfonden, the European Foundation for the Study of Diabetes, the NovoNordisk Foundation, Excellence of Diabetes Research in Sweden (EXODIAB), and the Family Ernfors-and OE&E Johanssons-Foundations.

## Supplementary Figures

**Fig. S1. Donor characteristics and validation of δ-cell markers in human islet preparations.**

A. Average HbA1c, BMI, and age (mean±SEM) for the studied donor sample populations, non-diabetic (ND; black) and type-2 diabetic (T2D; red) groups; dots indicate individual donors.

B. Immunocytochemical validation of δ-cell identification. Confocal images of dispersed human islet cells transduced with ad-Psst-NPY-EGFP (green). Cells were co-stained with antibodies against somatostatin (left, red), glucagon (middle, red), or insulin (right, red). EGFP-positive cells show strong overlap with somatostatin, confirming δ-cell identity, and no co-localization with glucagon or insulin. Scale bar: 50 µm.

**Fig. S2. Docked granules in δ-cells.**

Average docked granule density (mean±SEM) in δ-cells from non-diabetic (ND) or type 2 diabetic (T2D) donors, and in control, with somatostatin (SST), insulin (INS), γ-aminobutyric acid (GABA), glucagon (GCG), exendin-4 (EXN), forskolin (FSK), or adrenaline (ADR), each tested at both 10 mM and 1 mM (with diazoxide). The indicated comparisons are significantly different (ANOVA, Tukey’s test).

**Fig. S3. Relationship between δ-cell exocytosis and donor age or BMI**

A. Correlation of K^+^-stimulated granule exocytosis (mean±SEM, with diazoxide) and donor body mass index (BMI, left) or age (right) in dispersed human δ-cells for 1 mM glucose (1G; light blue) and 10 mM glucose (10G; dark blue) in presence of somatostatin (SST 400 nM). Each point represents the average exocytotic response of a single donor.

B. As in A, but in presence of ADR (adrenaline, 5 µM),

C. As in A, but in presence of GCG (glucagon, 10 nM).

**Fig S4. Effects of inhibitory and excitatory signals on δ-cell electrical activity.**

Average membrane potential in dispersed ND δ-cells (3 mM glucose) during application of somatostatin (SST, 25 cells, 5 donors), adrenaline (ADR, 13 cells, 4 donors), or GABA (40 cells, 5 donors). Individual recordings were aligned to the time of treatment and averaged.

---

**Supplemental Table 1.** Number of cells and donors used, non-diabetic.

| Treatment | Condition | Donors | Cells.per.donor | Mean.of.cells.per.donor | Total.cells |
| --- | --- | --- | --- | --- | --- |
| GLU | 1G | 7 | 4 - 7 | 5.30 | 37 |
| GLU | 10G | 11 | 4 - 13 | 7.60 | 84 |
| ADR | 1G | 5 | 5 - 9 | 6.20 | 31 |
| ADR | 10G | 7 | 4 - 6 | 5.10 | 36 |
| EXN | 1G | 4 | 4 - 5 | 4.80 | 19 |
| EXN | 10G | 4 | 4 - 5 | 4.50 | 18 |
| FSK | 1G | 3 | 5 - 6 | 5.30 | 16 |
| FSK | 10G | 4 | 2 - 11 | 5.50 | 22 |
| GABA | 1G | 3 | 4 - 5 | 4.70 | 14 |
| GABA | 10G | 4 | 4 - 7 | 5.20 | 21 |
| GCG | 1G | 4 | 5 - 8 | 6.50 | 26 |
| GCG | 10G | 5 | 5 - 7 | 5.60 | 28 |
| INS | 1G | 4 | 4 - 8 | 5.80 | 23 |
| INS | 10G | 4 | 5 - 8 | 6.00 | 24 |
| SST | 1G | 4 | 5 - 6 | 5.20 | 21 |
| SST | 10G | 8 | 3 -13 | 6.90 | 55 |

*Note.* Treatment abbreviations: 1G (1mM glucose), 10G (10mM glucose), GLU (glucose), ADR (adrenaline), EXN (exendin-4), FSK (forskolin), GABA (GABA), GCG (glucagon), INS (insulin), SST (somatostatin).

**Supplemental Table 2.**
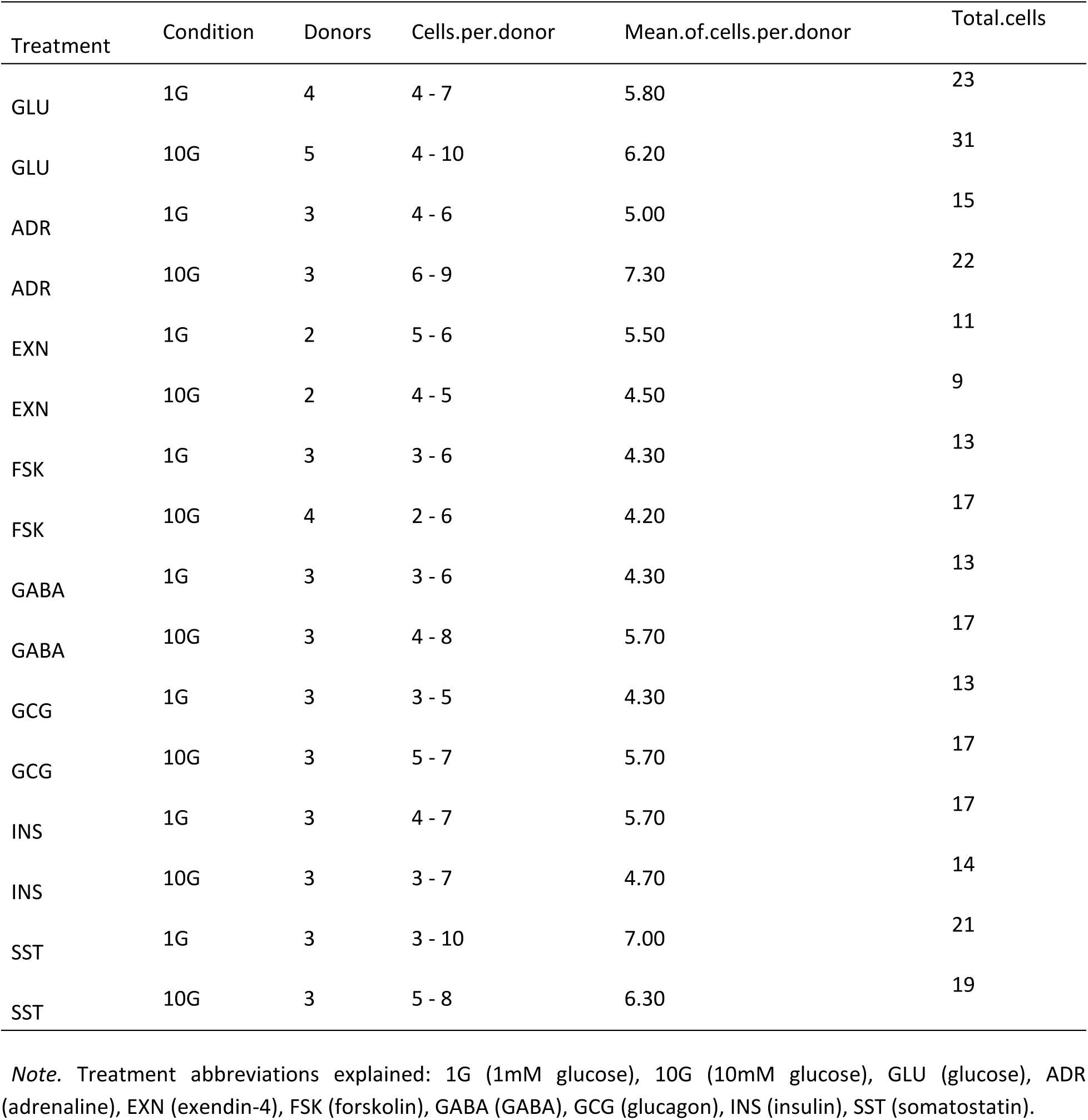
Number of cells and donors used, diabetic.

| Treatment | Condition | Donors | Cells.per.donor | Mean.of.cells.per.donor | Total.cells |
| --- | --- | --- | --- | --- | --- |
| GLU | 1G | 4 | 4 - 7 | 5.80 | 23 |
| GLU | 10G | 5 | 4 - 10 | 6.20 | 31 |
| ADR | 1G | 3 | 4 - 6 | 5.00 | 15 |
| ADR | 10G | 3 | 6 - 9 | 7.30 | 22 |
| EXN | 1G | 2 | 5 - 6 | 5.50 | 11 |
| EXN | 10G | 2 | 4 - 5 | 4.50 | 9 |
| FSK | 1G | 3 | 3 - 6 | 4.30 | 13 |
| FSK | 10G | 4 | 2 - 6 | 4.20 | 17 |
| GABA | 1G | 3 | 3 - 6 | 4.30 | 13 |
| GABA | 10G | 3 | 4 - 8 | 5.70 | 17 |
| GCG | 1G | 3 | 3 - 5 | 4.30 | 13 |
| GCG | 10G | 3 | 5 - 7 | 5.70 | 17 |
| INS | 1G | 3 | 4 - 7 | 5.70 | 17 |
| INS | 10G | 3 | 3 - 7 | 4.70 | 14 |
| SST | 1G | 3 | 3 - 10 | 7.00 | 21 |
| SST | 10G | 3 | 5 - 8 | 6.30 | 19 |

*Note.* Treatment abbreviations explained: 1G (1mM glucose), 10G (10mM glucose), GLU (glucose), ADR (adrenaline), EXN (exendin-4), FSK (forskolin), GABA (GABA), GCG (glucagon), INS (insulin), SST (somatostatin).

**Supplemental Table 3.** Donor characteristics.

| ND donors |  |  |  |  | T2D donors |  |  |  |  |
| --- | --- | --- | --- | --- | --- | --- | --- | --- | --- |
| ID | State | Age | BMI | HbA1c | ID | State | Age | BMI | HbA1c |
| 1414 | ND | 67 | 23,9 | 5,3 | 1105 | T2D | 51 | 26,1 | 7,8 |
| 1484 | ND | 60 | 24,2 | 5,2 | 1857 | T2D | 66 | 35,1 | 7,6 |
| 1780 | ND | 62 | 34 | 5,6 | 1864 | T2D | 65 | 33,3 | 7,3 |
| 1784 | ND | 70 | 23,3 | 5,4 | 1923 | T2D | 52 | 33 | 8,1 |
| 1882 | ND | 72 | 33,3 | 5,7 | 2034 | T2D | 64 | 29,6 | 7,7 |
| 1921 | ND | 55 | 23,8 | 5,4 | 2061 | T2D | 66 | 31,2 | 7 |
| 2108 | ND | 68 | 22,6 | 5,3 | 2211 | T2D | 63 | 37 | 7,9 |
| 2309 | ND | 54 | 42,6 | 5,5 | 2438 | T2D | 78 | 22,9 | 8,1 |
| 2340 | ND | 54 | 33,7 | 5,8 | 2536 | T2D | 55 | 35,1 | 6,5 |
| 2445 | ND | 62 | 22,5 | 4,8 | 2540 | T2D | 70 | 28,7 | 5,5 |
| 2516 | ND | 55 | 22,4 | 5,4 | 2471 | T2D | 77 | 26,3 | 5,8 |
| 2539 | ND | 75 | 23,8 | 5,2 | 2502 | T2D | 79 | 26,2 | 6 |
| 2550 | ND | 72 | 24,5 | 5,7 | 2509 | T2D | 69 | 31,7 | 6,7 |
| 2555 | ND | 58 | 21,8 | 5,4 | 2611 | T2D | 51 |  | 7,5 |
| 2585 | ND | 58 | 32,8 |  | 2618 | T2D | 80 | 27,7 | 6,3 |
| 2587 | ND | 40 | 44,4 | 6 | 2636 | T2D | 55 | 35,1 | 6,1 |
| 2595 | ND | 67 | 29 | 5,2 | R347 | T2D | 57 | 27,9 | 6,3 |
| 2606 | ND | 46 | 19 | 5,3 | R470 | T2D | 56 | 32,6 | 6,6 |
| 2608 | ND | 49 | 27,5 | 5,2 |  |  |  |  |  |
| 2673 | ND | 72 | 23,4 | 5,4 |  | mean | 64,1 | 30,6 | 6,93 |
| 2681 | ND | 59 | 35,2 | 5,5 |  | SEM | 2,3 | 1,0 | 0,20 |
| 2489 | ND | 65 | 30,1 | 6 |  |  |  |  |  |
| 2491 | ND | 71 | 22,4 | 5,9 |  |  |  |  |  |
| 2494 | ND | 81 | 21,1 | 5,3 |  |  |  |  |  |
| 2496 | ND | 73 | 23,2 | 5,6 |  |  |  |  |  |
| 2498 | ND | 36 | 18,9 | 5,1 |  |  |  |  |  |
| 2501 | ND | 53 | 26,6 | 5,1 |  |  |  |  |  |
| 2508 | ND | 70 | 22 | 5,1 |  |  |  |  |  |
| 2510 | ND | 52 | 26,9 | 5,5 |  |  |  |  |  |
| 2512 | ND | 56 | 24,6 | 5,8 |  |  |  |  |  |
| 2516 | ND | 55 | 22,4 | 5,4 |  |  |  |  |  |
| 2517 | ND | 51 | 28,1 | 5,4 |  |  |  |  |  |
| 2525 | ND | 63 | 22,1 | 5,3 |  |  |  |  |  |
| 2527 | ND | 43 | 18,9 | 5,2 |  |  |  |  |  |
| 2546 | ND | 61 | 24,5 | 5,4 |  |  |  |  |  |
| 2547 | ND | 62 | 29,4 | 5,6 |  |  |  |  |  |
| 2573 | ND | 55 | 32,1 | 5,6 |  |  |  |  |  |
| 2583 | ND | 49 | 23,6 | 5,2 |  |  |  |  |  |
| 2599 | ND | 54 | 18 | 5,4 |  |  |  |  |  |
| 2603 | ND | 69 | 31,1 | 5,8 |  |  |  |  |  |
| 2610 | ND | 62 | 22,2 | 5,5 |  |  |  |  |  |
| R385 | ND | 47 | 31 | 5,1 |  |  |  |  |  |
| R442 | ND | 53 | 12,8 | 5,7 |  |  |  |  |  |
| R536 | ND | 37 | 28,2 | 5,2 |  |  |  |  |  |
| R538 | ND | 61 | 32,9 | 5,5 |  |  |  |  |  |
|  | mean | 59,0 | 26,2 | 5,43 |  |  |  |  |  |
|  | SEM | 1,5 | 0,9 | 0,04 |  |  |  |  |  |

