## Supplementary figures and images for "Pancreatic δ-cells are resistant to auto- and paracrine inhibition in human type-2 diabetes"

### Supplementary figures

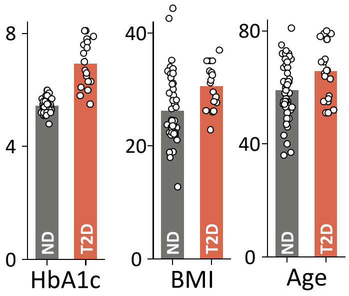

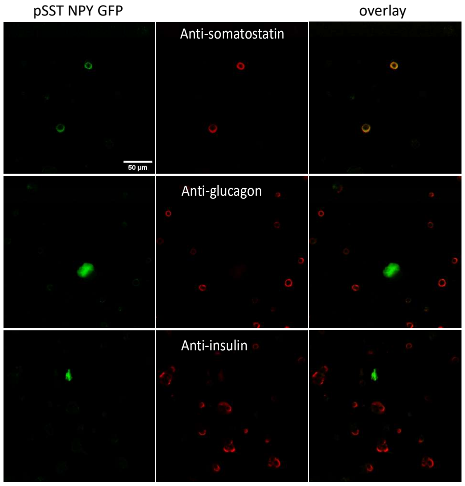

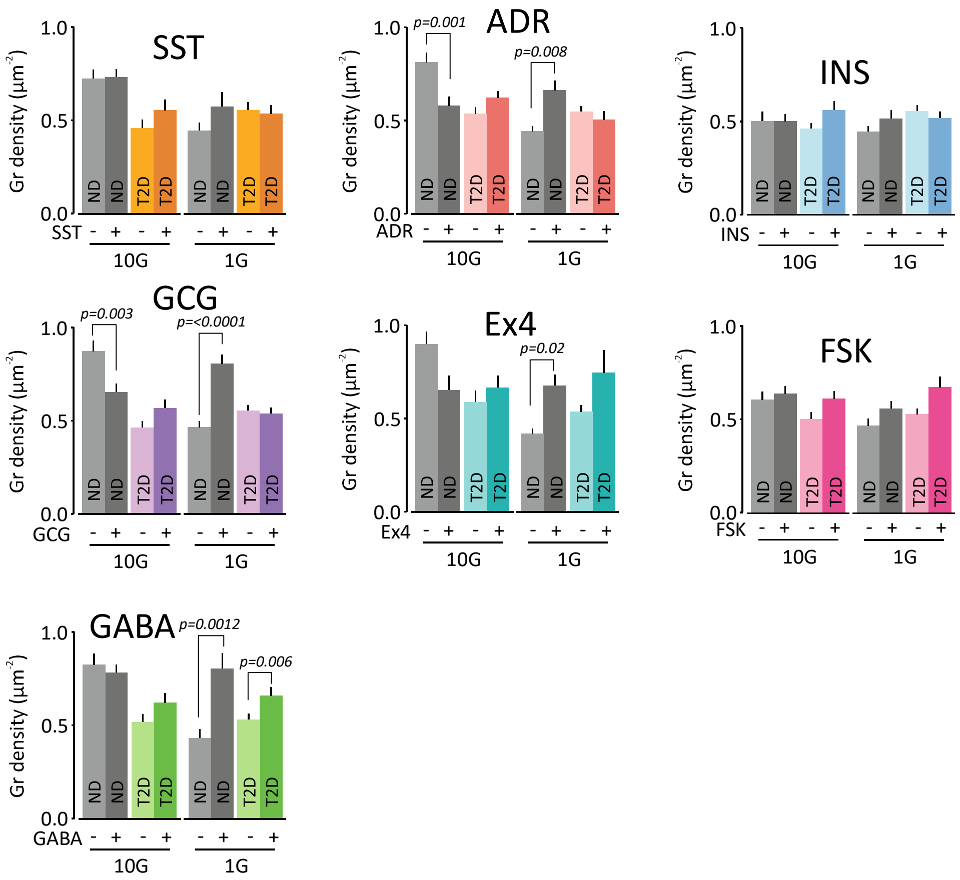

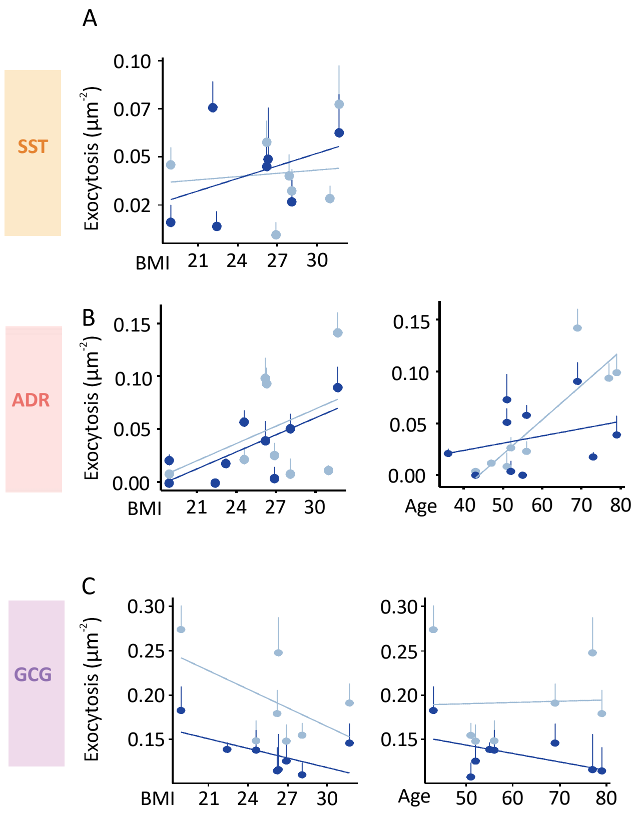

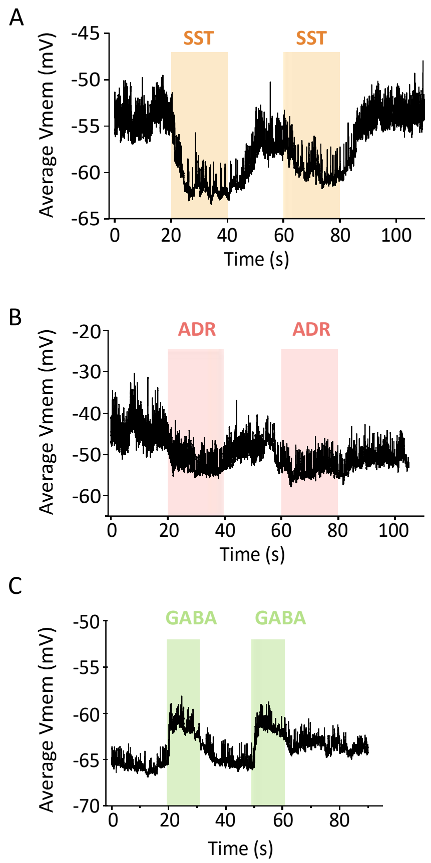
